# Possibilities of skin coat color-dependent risks and risk factors of squamous cell carcinoma and deafness of domestic cats inferred via RNA-seq data

**DOI:** 10.1101/2023.07.17.548980

**Authors:** Akinori Awazu, Daigo Takemoto, Kaichi Watanabe, Naoaki Sakamoto

## Abstract

The transcriptome data of skin cells from domestic cats with brown, orange, and white coats were analyzed using a public database to investigate the possible relationship between coat color-related gene expression and squamous cell carcinoma risk, as well as the mechanism of deafness in white cats. We found that the ratio of the expression level of genes suppressing squamous cell carcinoma to that of genes promoting squamous cell carcinoma might be considerably lower than the theoretical estimation in skin cells with orange and white coats in white-spotted cat. We also found the possibility of the frequent production of KIT lacking the first exon (d1KIT) in skin cells with white coats, and d1KIT production exhibited a substantial negative correlation with the expression of SOX10, which is essential for melanocyte formation and adjustment of hearing function. Additionally, the production of d1KIT was expected to be due to the insulating activity of the feline endogenous retrovirus 1 (FERV1) LTR in the 1st intron of *KIT* and its CTCF binding sequence repeat. These results contribute to basic veterinary research to understand the relationship between cat skin coat and disease risk, as well as the underlying mechanism.

## Introduction

Domestic cats are among the most important companion animals, and feline health occupies an important position in human society. It has also recently been suggested that dementia and breast cancer in cats share various similarities with Alzheimer’s disease (Chambers et al., 2015; Sordo et al., 2021) and triple-negative breast cancer in humans (Wiese et al., 2013; Caliari et al., 2014; Cannon, 2015; Thomas, 2015; Adega et al., 2016; Howard et al., 2020; Sommerville et al., 2022), respectively. Therefore, advances in feline medicine are expected to make important contributions to advances in human medicine.

In recent years, research on various feline diseases has progressed, leading to the development of vaccines for various lethal infectious diseases caused by virus such as Feline leukemia virus (FeLV) (Hofmann-Lehmann et al., 2020) and Feline immunodeficiency virus (FIV) (Westman et al., 2019; Andrade et al., 2022), therapeutic drugs for Feline infectious peritonitis (FIP) (Pedersen et al., 2019; Cook et al., 2022), and drugs for kidney disease (Sugisawa et al., 2016; Arai et al., 2016) that have been put into practical use, consequently extending the lifespan of domestic cats. However, these methods have only solved some diseases in cats. Basic medical research focusing on the genetics and physiology of cats is essential for establishing preventive and therapeutic methods for various feline diseases.

Domestic cats have different colors and patterns on their skin and coat, which vary from individual to individual depending on the combination of genes related to color (white, orange, brown, black, etc.) and patterns (agouti, tabby, dilute, etc.) (Lyons et al., 2005; Schmidt-Küntzel et al., 2005; Imes et al., 2006; Ishida et al., 2006; Schmidt-Küntzel et al., 2009; Eizirik et al., 2010; David et al., 2014; Lyons, 2021), and the pattern of X-chromosome inactivation in female. In addition, certain skin colors in cats have been suggested to be associated with specific diseases. For example, light-skinned individuals have been suggested to have a higher risk of squamous cell carcinoma than dark-skinned individuals (Dorn et al., 1971; Thomson, 2007; Murphy, 2013). This tendency has been attributed to the fact that light-colored individuals with less melanin have a weak resistance to ultraviolet (UV) rays. However, squamous cell carcinoma also develops in the lining of the oral cavity, tongue, and tonsils, which are less likely to be directly exposed to UV light. Therefore, there may be other marked factors besides UV light (Baer et al., 1993; Bertone et al., 2003; Sabattini et al., 2010; Munday et al., 2011; Munday et al., 2012; Dermatology et al., 2016, Sequeira et al., 2022).

Individuals with a white coat all over their bodies develop deafness at a high rate (Bergsma et al., 1971; Strain, 1996; Geigy et al., 2007; Ryugo et al., 2012; Strain, 2015; Kral et al., 2015; Strain, 2017; Mari et al., 2019; Bianchi et al., 2020). This disease is thought to result from the inhibition of melanin synthesis and the differentiation of melanocytes form the neural crest cells when *White* allele is present at white locus (Geigy et al., 2007; David et al., 2014; Strain, 2017). In a cat with white coat, a sequence derived from feline endogenous retrovirus 1 (*FERV1*) is inserted at the white locus in the first intron of the tyrosine protein kinase (*KIT*) gene (David et al., 2014). The *white spotting* allele of an individual with white-coated skin on a part of the body surface (white-spotted cat) contains the full-length sequence of *FERV1* (7125 bp). In contrast, the *White* allele of an individual whose whole body is completely covered with a white coat (dominant white cat) contains only the long terminal repeat (LTR) of *FERV1* (622 bp) and not the full length. This suggests that the presence of LTR caused a decrease in melanin synthesis. However, the role of LTR and the mechanism through which melanin synthesis is suppressed remains unclear. White coat skin formation with decreased melanin production is also associated with deafness in other animals, such as dogs, miniature pigs, and yaks (Strain, 2015; Strain, 2017; Qi et al., 2022). However, the underlying mechanisms have not been elucidated.

Thus, the relationship between the mechanisms that determine skin coat color and disease risk remains poorly understood. Therefore, in this study, we analyzed the RNA-seq data obtained from differently colored domestic cat skin cells published in a public database. Through this analysis, the possible relationship between skin color and squamous cell carcinoma risk and deafness as well as the mechanism by which genes that determine skin coat color is involved in these diseases were inferred.

This study had two further aims. First was to report the relationship between coat color and disease risk in domestic cats and its mechanism. The second was to promote basic veterinary research on feline diseases and data accumulation that serves as backbone for such research. Unfortunately, the availability of experimental data for domestic cats is currently substantially limited compared to that for humans and canines. In this study, only one replicate per skin cell color was available for RNA-seq analysis, which limited the application of conventional methods to assess differentially expressed genes (DEGs). Therefore, only forecast scenarios based on limited analysis could be determined. However, this study is expected to achieve both purposes by providing the motivation for various experimental studies and additional data analysis to validate the present hypothesis.

## Materials and Methods

### Quantification of transcriptome data in 3 types of skin cells of *Felis catus*

From the Ensembl public database, specifically the file transfer protocol (ftp) site, mapped RNA-seq data on the *Felis catus* genome, Fca126_mat1.0, were obtained in bam and bam.bai formats. Data were retrieved from the orange skin cells of a cat with orange and white coats (referred to as cell O), white skin cells of the cat with orange and white coats (referred to as cell W), and skin cells of a cat ticked with white torties (referred to as cell T). Each cell line had one replicate and the original Sequence Read Archive (SRA) identifiers were SRS1335428, SRS1319391, and SRS1319719. Here, O and W cells were obtained from the same cat, whereas T cells were obtained from a different cat (https://www.ebi.ac.uk/biosamples/samples/SAMN04498546; https://www.ebi.ac.uk/biosamples/samples/SAMN04498547; https://www.ebi.ac.uk/biosamples/samples/SAMN04498539).

We should note the following facts for the samples:

1. Both cats belonged to the white-spotted cat classification, which means that the cell genomes contained the full-length sequence derived from *FERV1* in the first intron of the *KIT* gene.
2. Most of the body surface of cats with white tortie-ticked tabby is covered with a dark-colored coat, such as brown or black. Therefore, for the analysis, it was assumed that cell T represented a typical dark-colored skin cell. As mentioned below, this assumption was supported by the expression levels of *tyrosinase* (*TYR*) and *tyrosinase related protein 1* (*TYRP1*) genes, which are located in the “color locus” and “brown locus” loci.
3. Cats with O and W cells are carriers of lipoprotein lipase deficiency.

The annotation file of the *F. catus* genome (Fca126_mat1.0) in the gene transfer format (GTF) was obtained from the National Centre for Biotechnology Information (NCBI) database (Bethesda, MD, USA). In the file, the chromosome numbers were converted from NC_058368.1, NC_058369.1, etc. to A1, A2, etc.

Using FeatureCounts with the options -M -O --fraction (Liao et al., 2014), the number of reads for each gene described in the GTF file was obtained from the bam and bam.bai formatted files for cells T, O, and W. From these counts, the transcript level of each gene was calculated by dividing the read count by the total exon length. The total transcript level was defined as the sum of all the transcript levels of the genes. Based on this result, the transcripts per million (TPM) of each gene were calculated as 1000000 * [transcript level] / [total transcript level] (see Table S1).

It is worth noting that the TPM values of the *TYR* and *TYRP1* genes in T cells were more than ten times higher than those in W cells (Table S1). This observation supports the previous assumption that T cells were sampled from dark-colored skin rather than white skin in cats with a ticked tabby and white torties.

### Size factor of RNA-seq data

As an example of size factors of RNA-seq data, the median of the ratios (MOR) defined by the recent literature for DESeq (Anders, et al, 2010) was estimated using read count data. For cells T, O, and W, the MORs were obtained 0.952545606, 1.201469023, and 0.868500362, respectively, and [standard deviation of MOR over three cells] / [average MOR over three cells] was obtained 0.140326793. Based on these results, it was expected that the size factor does not considerably affect the comparison of TPM values.

### Quantification of exon transcription levels of *KIT* in 3 types of skin cells of *F. catus*

A GTF file specifying each exon region of *KIT* was obtained from the GTF file Fca126_mat1.0. The read count in each exon region of *KIT* was obtained from bam and bam.bai formatted files with FeatureCounts (using the options: -t feature -M -O--fraction) (Liao et al., 2014) for T, O, and W cells. The transcript level of each region was calculated as the ratio of read count to region length. From the results, the TPM in each exon region was obtained as 1000000 * [transcript level] / [total transcript level] (Table S2), where the total transcript level uses the value calculated in the previous section.

### Extraction of skin coat color-dependent differentially expressed genes (SCDEGs)

The median TPM values (Table S1) obtained for all genes in all the cells (T, O, and W) were approximately 0.987. Based on this fact, we considered genes that met the following conditions as skin coat color-dependent differentially expressed genes (SCDEGs):

- The maximum TPM values obtained for T, O, and W were more than *u*-times of the minimum value.
- The maximum TPM values obtained for T, O, and W were *v* or higher.

The following arguments, *u* = 2 and *v* = 2 (approximately equal to twice the median value of TPM values) were assumed to estimate SCDEGs. Additionally, the SRY-Box Transcription Factor 10 (*SOX10*) was included as an SCDEG. Although the maximum expression level obtained from T, O, and W was 1.9 times the minimum value, which does not satisfy the above requirement, *SOX10* is one of the predominant factors in determining skin coat color (Mort et al., 2015; Strain, 2015; Qi et al., 2022).

Furthermore, the maximum expression of *SOX10* was nearly twice the minimum value. Therefore, *SOX10* was specifically considered an SCDEG (Table S3).

### Enrichment analysis

Pathway enrichment analysis of the SCDEGs was conducted using g:Profiler (https://biit.cs.ut.ee/gprofiler/gost) (Raudvere et al., 2019). For this analysis, “felis catus” was chosen as the target organism, and “only annotated genes” were selected as the target gene set. The focus of the analysis was to identify significantly enriched Kyoto Encyclopedia of Genes and Genomes (KEGG) pathways with a false discovery rate (FDR) threshold of less than 0.1.

### Classification into squamous cell carcinoma-promoting and suppressing genes

First, SCDEGs associated with the KEGG pathway were identified from the results obtained through the aforementioned pathway enrichment analysis. Next, using databases and prior literature, the extracted SCDEGs were categorized into the following groups:

1. Carcinoma-promoting genes: These genes exhibited an excessive increase in expression leading to promotion of squamous cell carcinoma.
2. Carcinoma suppressor genes: These were expressed at decreased levels or were mutated; these genes contributed to the development of squamous cell carcinoma.
3. Other genes: SCDEGs that did not fall into the above categories. Categorization was based on the information obtained from databases and previous studies (Table S5).

### Theoretical ratio of the number of “tumor-promoting genes” and “tumor-suppressor genes”

A file named cancermine_collated.tsv, which contained cancer gene roles and citations, was obtained from the data download page (https://zenodo.org/record/7689627) of the public database CancerMine (http://bionlp.bcgsc.ca/cancermine/) (Lever et al., 2019). This file provided a list of oncogenes and tumor suppressor genes for each of the 526 cancer types covered in the database. The total number of oncogenes and tumor suppressor genes in all carcinomas was 18,994 and 11,295, respectively. Therefore, we assumed that the “theoretical ratio for all carcinomas” between the number of “carcinoma-promoting genes” and the number of “carcinoma-suppressor genes” could be estimated based on these data.

Additionally, CancerMine contains data of 17 types of squamous cell carcinomas (Table S4), where the total number of oncogenes and tumor suppressor genes in all squamous cell carcinomas was 904 and 584, respectively. Therefore, we also assumed that the “theoretical ratio for all squamous cell carcinomas” between the number of “carcinoma-promoting genes” and the number of “carcinoma-suppressor genes” could be estimated.

Although the database data used for estimating this theoretical ratio are specific to human cancer, it is reasonable to assume that the ratio of the number of genes would be similar between the two species because humans and cats are both mammals.

### Evaluation of skin coat color dependent squamous cell carcinoma risk

To assess the cancer risk, the expression levels of carcinoma-promoting and carcinoma-suppressor genes within the SCDEGs, specifically those contributing to pathways related to squamous cell carcinoma, were compared among T, O, and W cells using the following criteria:

1. The number of carcinoma-promoting genes whose expression level in cell X (X = T, O, or W) was at least twice the minimum expression level among cells T, O, or W was denoted as P_X_.
2. The number of carcinoma-suppressor genes whose expression level in cell X (X = T, O, or W) was at least twice the minimum expression level in cells T, O, or W was denoted as S_X_.
3. If P_X_ was greater than S_X_, a comparison was made with the theoretical ratio obtained earlier using an exact test of goodness-of-fit. If the resulting *p*-value is less than 0.05, it indicates that cell X is in a state where squamous cell carcinoma is more likely to develop and progress than in the other cells.

Now, it should be noted that whether the decreases in the expression levels of tumor suppressor genes in normal cells could be useful for predicting carcinogenesis is not trivial. Generally, mutations in some genes trigger tumorigenesis. For some carcinomas, however, recent studies suggest that candidate tumor suppressor genes, such as CDH1, PTEN, and KiSS1, expression levels decreased by epigenetic silencing, rather than by deletion, cause carcinogenesis (Wang, et al., 2019). Thus, the ratio of excessively expressed carcinoma promoting and suppressor genes in normal cells were focused because such ratio might provide nonnegligible contributions to squamous cell carcinoma risk.

### Evaluations of motif sequence of CTCF binding and promoter activity in white locus LTR

T, O, and W cells were derived from individual cats partially surrounded by white skin and fur. In the genomes of these white-spotted cats, an LTR sequence derived from *FERV1* was located in the first intron of *KIT*. To determine the specificity of this LTR sequence, we searched for binding motif sequences of the CCCTC-binding factor (CTCF) and motif sequence for core promoter elements within the LTR.

For this purpose, we utilized the CTCFBS Prediction Tool available in CTCFBSDB 2.0 (https://insulatordb.uthsc.edu/) (Ziebarth et al., 2013) and Eukaryotic Core Promoter Predictor (YAPP: http://www.bioinformatics.org/yapp/cgi-bin/yapp.cgi). The former tool is based on a database containing the CTCF-binding motif sequences from humans and mice, where the given sequence is considered to involve CTCF-binding activity if the score of this tool is larger than 3. The latter tool is based on the statistical analysis of data sets of eukaryotic core promoter elements (Cartharius et al., 2005; Gershenzon et al., 2005; Jin et al., 2006), where the sequence with a score larger than 0.8 (0 < score < 1) is considered to exhibit promoter activity; the sequences with a score equal to or larger than 0.9 were focused in this study.

Feline CTCF has more than 99% of the same amino acid sequence as human CTCF, and all of the DNA-binding motif amino acid sequences are the same (Fig. S1). Therefore, although the CTCFBSDB database was primarily constructed based on research conducted in humans and mice CTCF, the use of this database is reasonable for the CTCF analysis of domestic cats.

### Statistical analysis

*P*-values for various statistical analyses were calculated using R (version 4.2.3).

## Result

### SCDEGs and results of KEGG pathway enrichment analysis

To investigate the relationship between SCDEGs (Table S3) and the risk of squamous cell carcinoma, KEGG pathway enrichment analysis was conducted using g:Profiler (Table S5a). The analysis identified the following pathways related to squamous cell carcinoma in which significantly larger number of SCDEGs were involved.

1. Pathways in cancer (FDR ∼ 3.58×10^-3^)
2. The p53 signaling pathway (FDR ∼ 0.013×10^-2^)
3. Human papillomavirus (HPV) infection (FDR ∼ 8.33 × 10^-2^)

HPV infection is implicated in the development of human squamous cell carcinoma and cervical cancer (Hennessey et al., 2009; Husain et al., 2017; Augustin et al., 2020; Bao et al., 2021; Chrysovergis et al., 2021; The et al., 2021; Karina, 2022; Kwon et al., 2022). Feline papillomavirus (FPV) type-2 has been proposed as the primary cause of squamous cell carcinoma in domestic cats (Munday et al., 2008; Munday et al., 2011; O’Neill et al., 2011; Munday et al., 2012; Thomson et al., 2016; Munday et al., 2017; Carrai et al., 2020; Yamashita-Kawanishi et al., 2021; Teh et al., 2021; Altamura et al., 2022). HPV and FPV are distinct viruses; however, their *E5*, *E6*, and *E7* oncogenes exhibit similar features (Jiang et al., 2014; Altamura et al., 2016; Karina, 2022). Additionally, the pathogenesis of FPV-2 infection share pathways with HPV-induced pathogenesis and carcinogenesis (Teh et al., 2021; Karina, 2022). Consequently, genes involved in processes driven by HPV infection should also be considered genes associated with squamous cell carcinoma.

KEGG pathway enrichment analysis might change depending on the chosen SCDEGs determined by *u* and *v* values used. Here, to clarify the limitation of the above result, the cases with *u* and *v* larger than 2 were exceptionally considered. Consequently, the abovementioned three KEGG pathways related to squamous cell carcinoma contained a substantial large number of SCDEGs (FDR < 0.1) when *u* ≤ 2.5 and *v* ≤ 4.5 (Table S5b).

Notably, the same analysis using RNA-seq read data of mesenteric lymph node cells from 4 different domestic short hair cats did not exhibited similar results, where “pathways in cancer” and “Human papillomavirus (HPV) infection” did not contain significantly larger number of SCDEGs (Supporting information 1, Table S6a, and S6b). This fact may support that the SCDEGs among T, O, and W cells and their squamous cell carcinoma-related characteristics were rather specific than emerged due to variations among individuals.

### Assessment of squamous cell carcinoma risk by comparing the expression of carcinoma-promoting and carcinoma-suppressing genes

SCDEGs that contributed to cancer pathways, the p53 signaling pathway, and human papillomavirus infection were classified into squamous cell carcinoma-promoting and squamous cell carcinoma-suppressing genes based on previous studies (Table 1, Table S7). For each of the T, O, and W cells, the P_X_ and S_X_ that exhibited stronger expression than the others were counted. The relationship between P_X_ and S_X_ (X = T, O, W) was compared to the theoretical ratio of the number of carcinoma-promoting and carcinoma-suppressor genes using the exact goodness of fit test (Table 2).

**Table 1:**
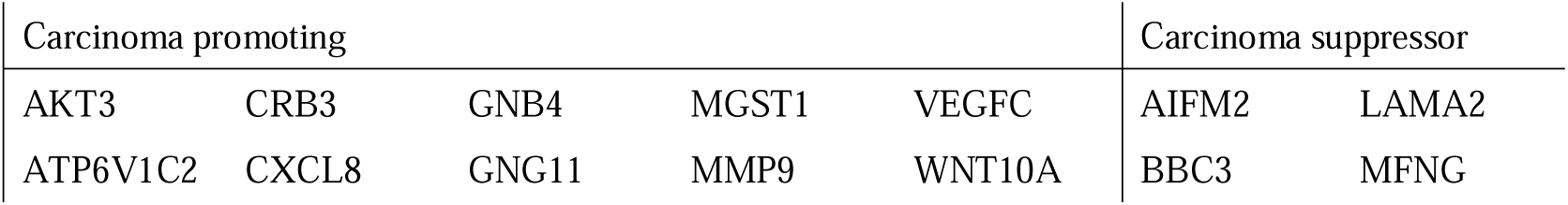
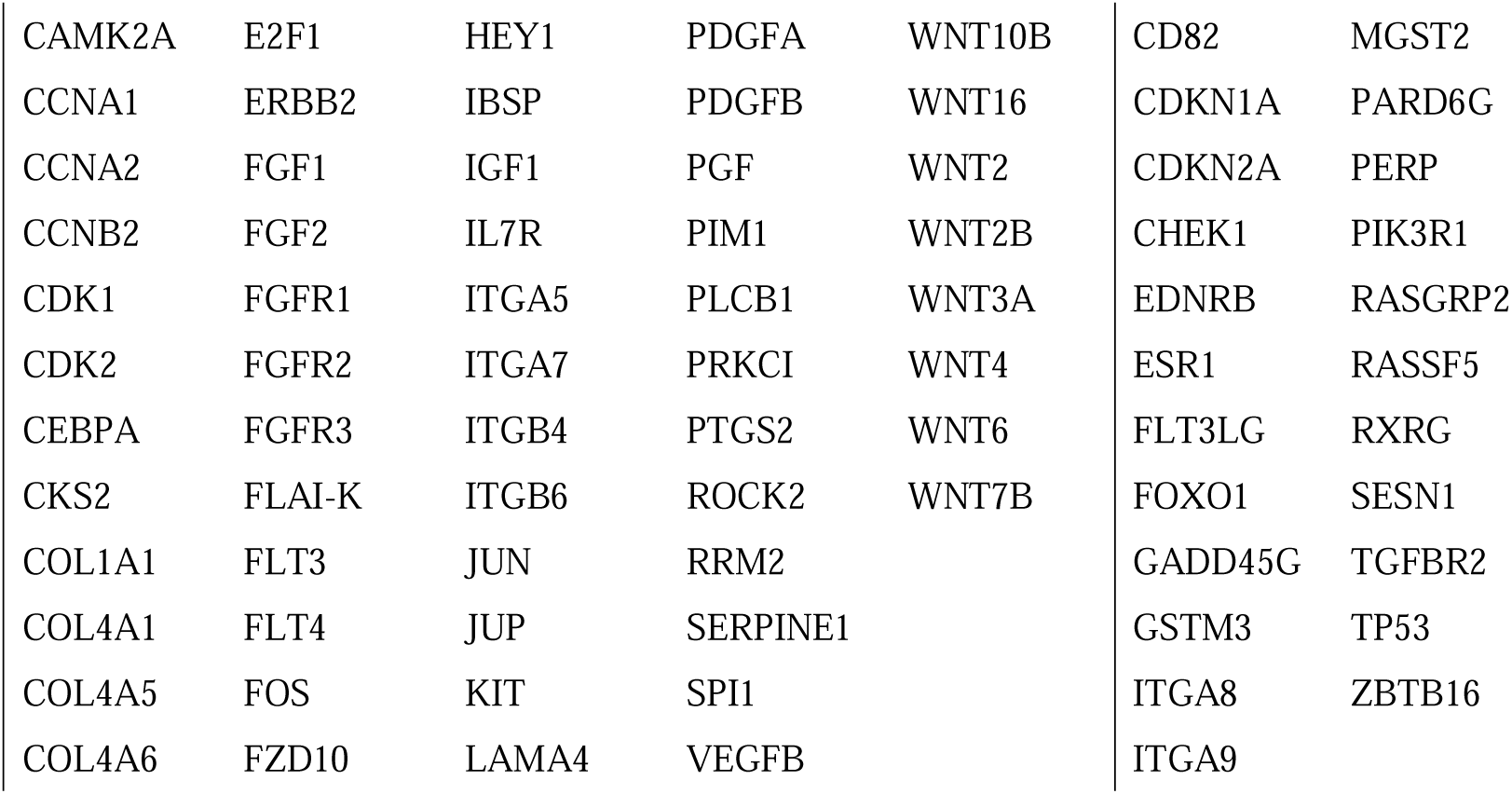
List of squamous cell carcinoma promoting and suppressor SCDEGs.

**Table 2:** Number of squamous cell carcinoma promoting and suppressor genes, which are more strongly expressed, for cells T (P_T_ and S_T_), O (P_O_ and S_O_), and W (P_W_ and S_W_), and *P*-values of exact test of goodness-of-fit between promoting-suppressor genes of each case and the theoretical ratio, where ALL indicate the theoretical ratio for all carcinomas and SCC indicate squamous cell carcinomas classified in CancerMine (Table S4). *P*-value of Fisher’s exact test for a 2X 3 table with P_T_, S_T_, P_O_, S_O_, P_W_, and S_W_ was obtained ∼ 0.1296.

|  | T | O | W | ALL | SCC |
| --- | --- | --- | --- | --- | --- |
| Carcinoma promoting genes | $P_T = 27$ | $P_O = 32$ | $P_W = 31$ | 18994 | 904 |
| Carcinoma suppressor genes | $S_T = 17$ | $S_O = 8$ | $S_W = 9$ | 11295 | 584 |
| $P$-value (for ALL) | 0.8768 | 0.02246 | 0.05194 | | |
| $P$-value (for SCC) | 1 | 0.01421 | 0.03445 | | |

The analysis revealed that the P_O_ in O cell was significantly higher than the number of carcinoma-suppressor genes (S_O_) (*P* = 0.02246 compared with the theoretical ratio for all carcinomas, and 0.01421 compared with all squamous cell carcinomas). The relationship between P_W_ and S_W_ displayed similar features, although the *P*-value obtained was slightly larger (*P* = 0.05194 compared with the theoretical ratio for all carcinomas, and 0.03445 compared with all squamous cell carcinomas). The values for P_T_, P_O_, and P_W_ were similar. In contrast, S_O_ and S_W_ values were lower than that of S_T_ (*P* value = 0.1296 using the Fisher’s exact test), indicating that the expression levels of carcinoma-suppressing genes were considerably lower in O and W cells than in T cells. A similar result was also confirmed when *u* = 2.5 and *v* = 4.5 (Table S5c).

### CTCF binding motif repeat in LTR at white locus in 1st intron of *KIT* gene

White-coated cats possessed a DNA sequence derived from *FERV1* located in the first intron of the *KIT* gene. Furthermore, both *White* and *white spotting* alleles includes a 5’ LTR region, which is expected to inhibit melanocyte differentiation (David et al., 2014). In this study, we examined the characteristics of the LTR sequence by evaluating the presence of CTCF-binding motifs in this region using CTCFBS Prediction Tool. The analysis identified putative seven CTCF-binding motifs within the LTR that showed similarity to known CTCF-binding sites (Fig. 1a). Furthermore, these motif sequences were arranged in a repeat pattern with a spacing of 46 base pairs between them.

**Figure 1:**
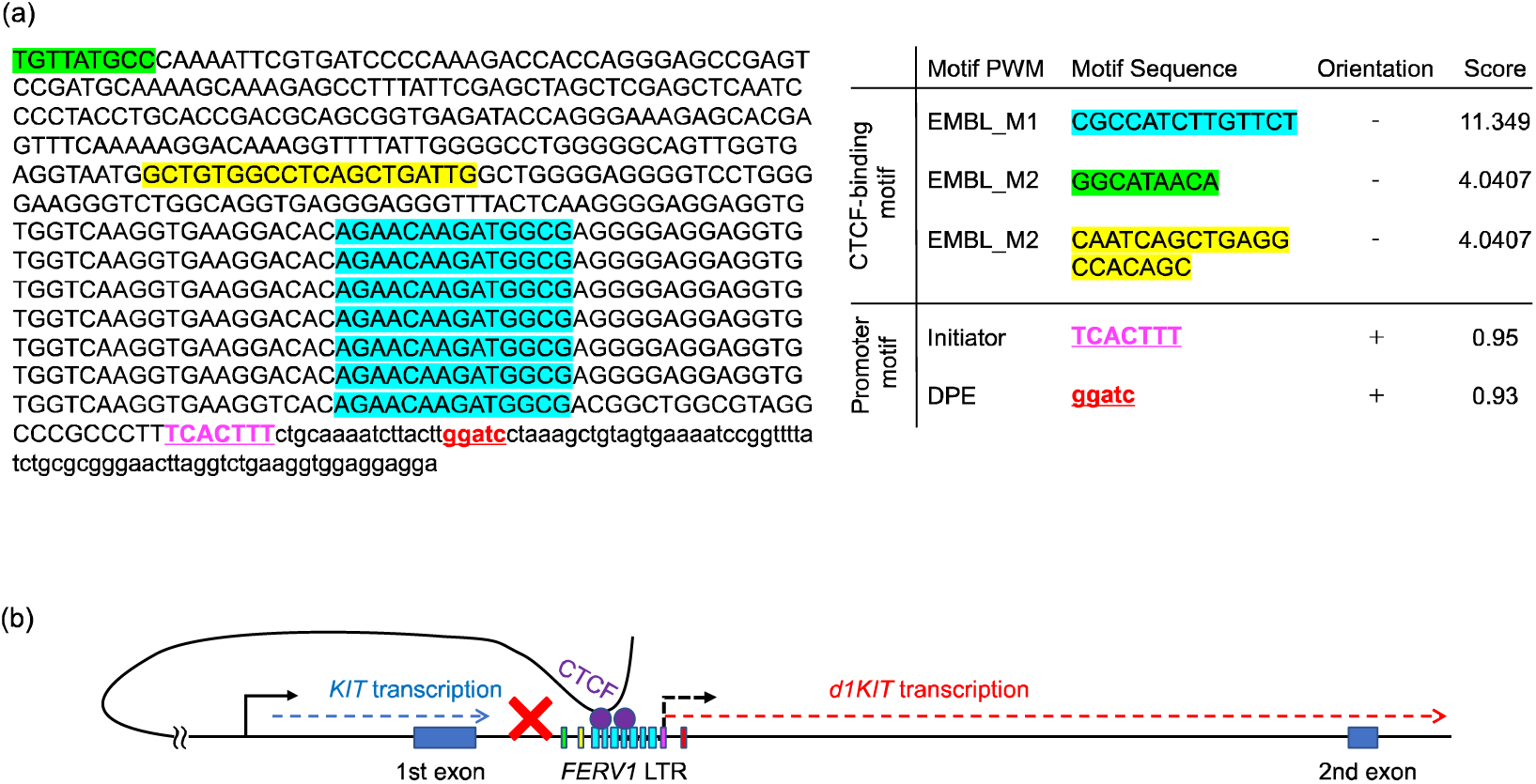
Positions of motif sequences with higher scores of CTCF-binding and core promoter activity in *FERV1* LTR in the first intron of the *KIT* gene. Detailing position and sequence of each CTCF-binding motif (cyan, green, and yellow shadowed) and each core promoter motif (magenta and red texts) (a), and illustration of inferred promoter and insulator activities of *FERV1* LTR (b). Capital letters indicated the sequence in LTR and lower case letters indicated outside of LTR.

### 1st exon-lacking KIT (d1KIT) expression level increased in white skin cell

The repeat region of the putative CTCF-binding motif within the LTR of the first *KIT* intron suggested a potential role of the LTR by binding of CTCF and functioning as an insulator, thereby acting as a boundary for topologically associated domains (TADs). *CTCF* has been found to be enriched at the edges of chromatin loops that form TADs and cooperates with cohesins to regulate TAD boundary formation (Parelho et al., 2008; Wendt et al., 2008; Merkenschlager et al., 2016). Genes located within neighboring TADs are generally known to be independently regulated in their expression.

Retroviral LTRs typically function as transcription promoters (Klaver et al., 1994; Pereira et al., 2000; Dunn et al., 2003; Cohen et al., 2009). The *FERV1* LTR may also possess promoter activity because the prediction tool YAPP found an initiator motif (INR), which is a typical consensus motif sequence of core promoter region, at the downstream of the repeat region of the putative CTCF-binding motif in LTR (Fig. 1a). Additionally, a downstream promoter element (DPE), which is also typical consensus motif sequence of core promoter region, was also found at the downstream of the INR (Fig. 1a). The DPE and the INR can function as TFIID-binding site cooperatively, leading to accurate and efficient transcription (Burke and Kadonaga, 1996, 1997). Therefore, if CTCF-binding motifs exhibit insulating activity, it is possible that the second exon and downstream regions of the *KIT* gene can be transcribed independently of the native promoter, utilizing the region regulated by above pair of INR and DPE as a core promoter (Fig. 1b). Now, we tentatively named the *KIT* transcribed in such a manner and lacking the first exon *d1KIT* and quantified its expression level in each cell type in the next section.

Based on the estimated TPM values provided in Tables S1 and S2, the ratio of [TPM of the 1st exon] / [TPM of KIT] in T and O cells was in the range of 0.8 and 0.84, respectively. Additionally, the ratio of [Average TPM of 2nd-22nd exons] / [TPM of KIT] in T, O, and W cells was close to 1. However, in W cell, the ratio of [TPM of 1st exon] / [TPM of KIT] was approximately 0.48, which was close to half of the values observed in the other cells (Fig. 2). This suggests that approximately half of the *KIT* mRNAs generated by W cells lacked the first exon.

**Figure 2:**
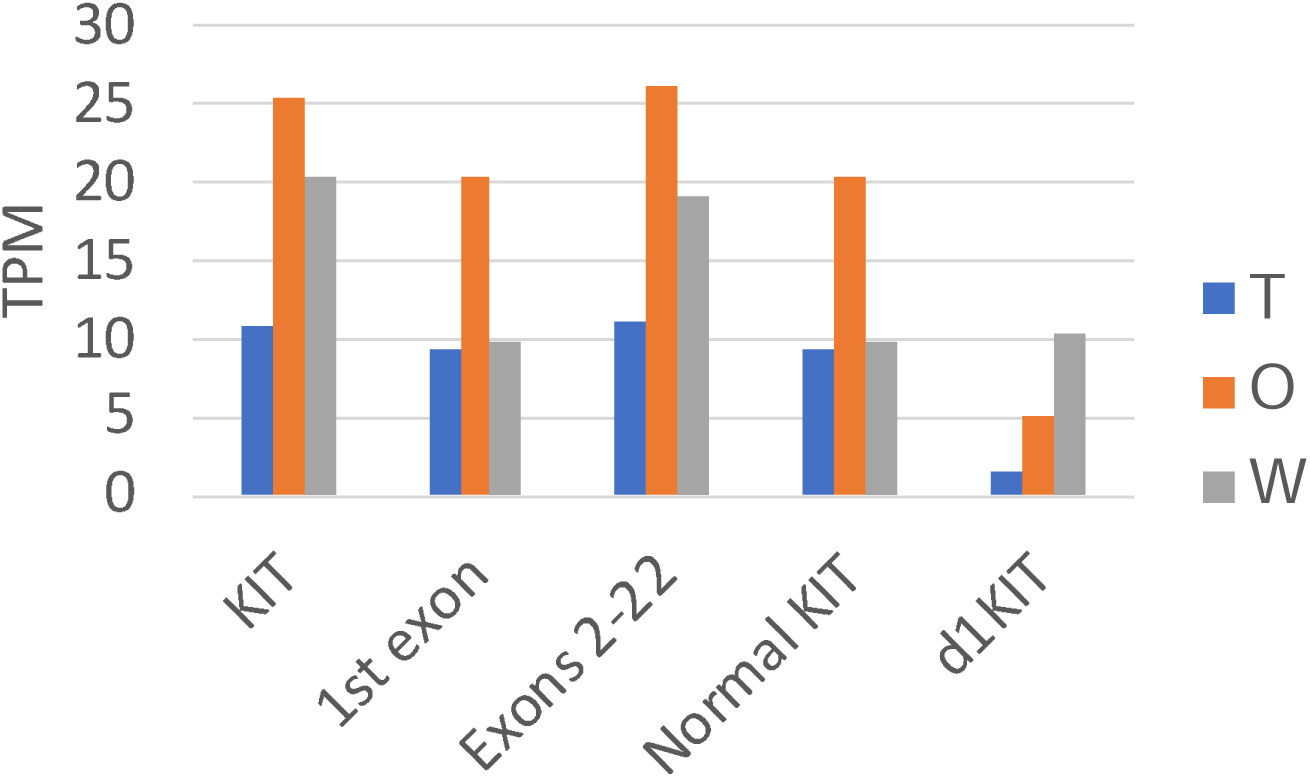
TPM of the following: entire KIT, 1^st^ exon of KIT, averages over 2^nd^ ∼ 22^nd^ exon, normal KIT, and d1KIT of T, O, and W cells. Values indicate TPM and length of each green bar is proportional to TPM.

The approximate transcript levels (TPM) of normal *KIT* and *d1KIT* in T, O, and W cells were estimated as follows. First, since the amount of normal *KIT* mRNA was expected to consist of those of the first exon and the second and subsequent exons, [TPM of normal KIT] was assumed to be the minimum value between [TPM of 1st exon] and [TPM of KIT]. Second, the TPM of d1KIT was estimated as follows: [TPM of d1KIT] = [TPM of KIT] – [TPM of normal KIT]. The TPM of normal *KIT* in T, O, and W cells was approximately 9.35, 20.50, and 9.87, respectively, whereas that of *d1KIT* in T, O, and W cells was approximately 1.67, 5.00, and 10.49, respectively (Fig. 2). Here, the ratio [TPM of d1KIT]/[TPM of normal KIT] in T, O, and W cells was 0.179929164, 0.244270964, and 1.063338179, respectively.

Notably, the transcript levels of normal KIT and d1KIT could not be obtained directly from short read RNA-seq data as analyzed in this study; however, they could be estimated using various methods. Thus, to confirm the validity of the abovementioned results, [d1KIT expression]/[Normal KIT expression] was estimated using the maximum likelihood estimate (MLE) method, using the number of read of each exon of KIT. Here, the [d1KIT expression]/[Normal KIT expression] for T, O, and W cells was 0.191814285, 0.259627163, and 1.112849284, respectively, which seemed similar to the above ratio [TPM of d1KIT]/[TPM of normal KIT] (Supporting information 2). Therefore, [TPM of normal KIT] and [TPM of d1KIT] were used as transcript levels of normal KIT and d1KIT in the following arguments.

### *d1KIT* expression correlated negatively to gene expressions for melanocyte formation

To determine how normal KIT and d1KIT are involved in melanocyte formation, we examined the correlation between their expression levels and those of typical genes involved in melanoblast and melanocyte formation (Table S8). The expression level of *d1KIT* showed a significant negative correlation with the expression level of *SOX10*, with a correlation coefficient of approximately −0.99957 (*P*-value ∼ 0.01876) (Fig. 3). Furthermore, strong negative correlations were observed with the expression levels of melanocyte-related *DKK4* and *TYRP1*, with correlation coefficients of approximately −0.98777 (*P*-value ∼ 0.09968) and −0.98000 (*P*-value ∼ 0.12763), respectively. In contrast, the correlation coefficient between the expression levels of normal *KIT* and melanocyte protein *PMEL* ∼ 0.94548 (*P*-value ∼ 0.20302) that exhibited the largest absolute value of correlation coefficient (Fig. 3). This fact suggested the expression level of normal KIT did not strongly correlate with the expression of genes involved in melanoblast and melanocyte formation.

**Figure 3:**
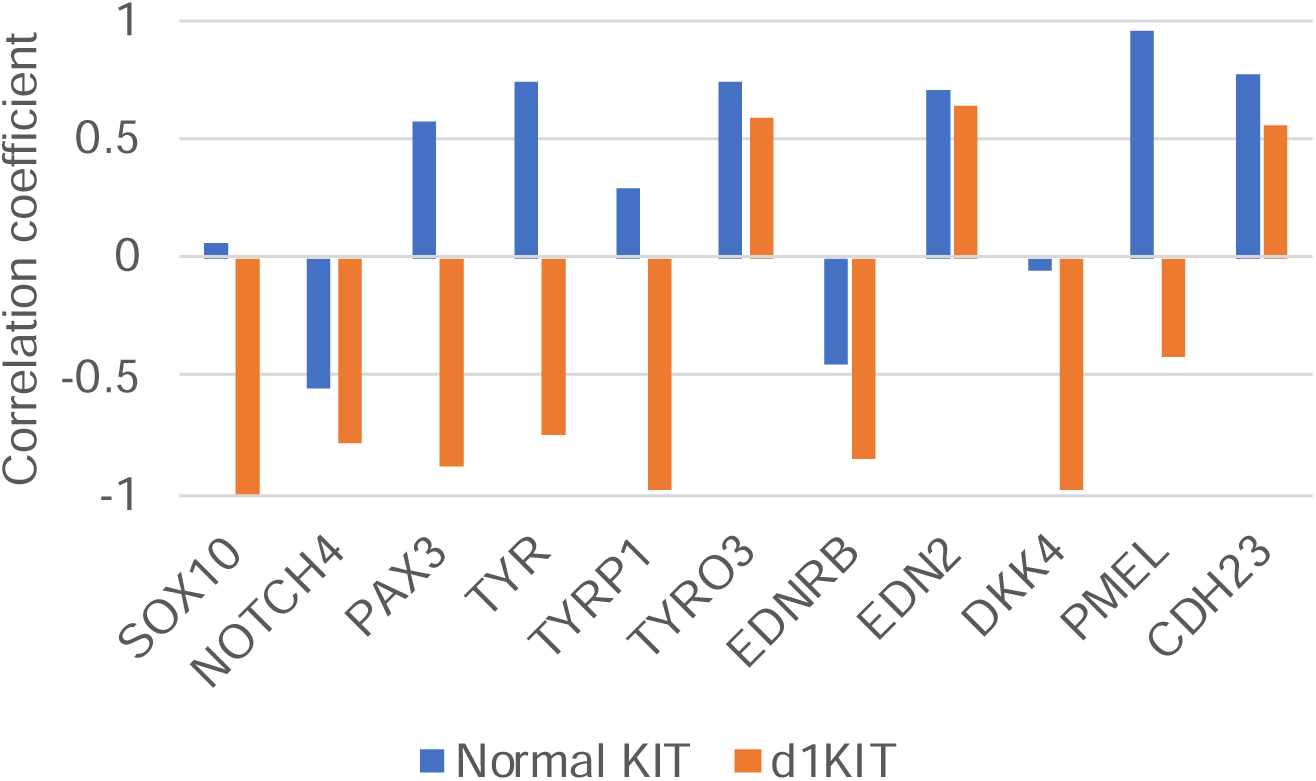
Correlation coefficients between TPMs of typical SCDEGs that are involved in melanoblast and melanocyte formation and TPM of normal KIT (Blue) or d1KIT (Orange). P-values of correlation coefficient between normal *KIT* and *PMEL* ∼ 0.20302, that between *d1KIT* and *SOX10* ∼ 0.01876, that between *d1KIT* and *Dkk4* ∼ 0.09968, and that between *d1KIT* and *TYRP1* ∼ 0.12763.

## Discussion

Squamous cell carcinoma tends to occur more frequently in light-coated cats, which may be due to a lack of melanin production, resulting in poor UV resistance. If this is the case, it would be expected that white-coated skin, without melanin production, would be most susceptible to carcinoma. However, in our analysis, we found the possibility that, in terms of gene expression, the expression level of a factor that suppresses squamous cell carcinoma was significantly lower in cells with orange-coated skin with *white spotting* allele. In other words, when considering gene expression, cells with not only white-but also orange-coated skin may have higher risks of cancer than cells with other skin colors.

We also observed that the ratio of the expression level of genes suppressing squamous cell carcinoma to that of genes promoting squamous cell carcinoma was lower than the theoretical estimation in cells with white coated skin with *white spotting* allele. It is worth noting that the orange- and white-coated skin cells analyzed in this study were sampled from the same individuals and were derived from carriers of lipoprotein lipase deficiency. These findings suggest two possible mechanisms underlying the risk of squamous cell carcinoma.

1. Cells for light-colored skin, such as white- and orange-coated skin, tend to exhibit low expression of carcinoma suppressor genes.
2. Cells derived from carriers with lipoprotein lipase deficiency tended to exhibit low expression of carcinoma-suppressor genes.

Validation of these possibilities through analysis of more accumulated data and additional experiments should be conducted in future.

Here, squamous cell carcinoma develops from cells in the epidermis. Thus, the ideal samples would be squamous cells or at least epidermal cells rather than skin cells as used in the present study. Feline squamous cell carcinoma is highly malignant. For example, oral squamous cell carcinoma has a poor prognosis with a one-year survival rate of less than 10% (Martin et al., 2011). However, there are no public RNA-seq data on squamous cells or epidermal cells derived from cats with different coat colors.

Therefore, in this study, the analysis of the data obtained from skin cells was performed to see the possibilities expected from it, knowing that there were several limitations. However, more accurate prediction of squamous cell carcinoma risk becomes possible by obtaining and analyzing the data through experiments of squamous cells and epidermal cells from cats with different coat colors. Such studies should be progressed in the future.

The contributions of each gene to carcinogenesis seemed to be different. For example, change in expression level of the *TP53* should be critical, while those in others might have moderate effects. However, there were no appropriate way to quantify the contribution weight of each gene to the carcinoma risks. Thus, in the present study, we assumed that such weight was approximately uniform as first trial analysis. However, the estimation of such weight is crucial for a more accurate carcinoma risk assessment.

In white-coated skin cells, a large amount of *d1KIT* mRNA, in which the 1st exon of the *KIT* gene is deleted, might be produced. Furthermore, we found the possibility that the LTR of *FERV1*, which was present in *White* and *white spotting* alleles at white locus in the first intron of the *KIT* gene in the genome of a domestic cat with white-coated skin, contains a repeated CTCF-binding site. This finding suggests the possibility that the transcription from the 1st exon and that from the 2nd exon are separately regulated by the presence of the promoter and insulator activities of this LTR, which provided the mechanism for d1KIT production.

Additionally, the amount of *d1KIT* transcript was found to exhibit a significant negative correlation with the expression levels of *SOX10*, as well as strong negative correlations with genes that induce differentiation into melanocytes. These results suggest the possibility that the insulating activity at the white locus, specifically the LTR of *FERV1*, resulted in the production of d1KIT and subsequent inhibition of the *SOX10* expression, which was crucial for melanocyte formation.

Notably, Sox10 mutations cause deafness in miniature pigs, leading to degeneration and a considerable decrease in the density of utricular hair cells, cochlear abnormalities, and saccular hypofunction (Qi et al., 2022). Therefore, when the aforementioned LTR activities occur throughout the body, it is conceivable that the entire skin becomes covered with white fur and deafness is induced. We hypothesized that this is the mechanism underlying dominant white cat deafness.

However, in white-spotted cats, from which the cells sampled in this study were derived, LTR activity and its influences on the KIT gene expression might be limited depending on cell lineages or cell types, since the full-length of FERV1 is present at *white* locus. For example, between O and W cells differences in 3D genomic structures of FERV1 other than LTR or those in DNA methylation status of the LTR promoter region might appear, and contributed to the around twice difference of expression of the d1KIT appeared. However, the precise mechanism remains unclear, and its elucidation is an issue for future research through the epigenetic measurements such as Hi-C and 4C-seq, methylation, and nucleosome binding around FERV1 in the first intron of KIT.

Furthermore, for both dominant white and white spotted cats, it was difficult to show a causal relationship between d1KIT production and *SOX10* expression only in the present study. Such relationship should be confirmed by future experiments.

As mentioned in the last part of introduction section, the availability of experimental data for *F. catus* (domestic cats) is currently significantly limited compared to that for humans and canines, which are also common companion animals. For instance, the number of *F. catus* RNA-seq samples registered in the public database COXPRESDB (https://coxpresdb.jp/RNA-seq) (Obayashi et al., 2023), which allows the search of co-expressed genes, is only 267, whereas the numbers for humans and canines are much larger (235,187 and 1,361, respectively). Additionally, the specific roles of each gene in the KEGG pathways obtained through enrichment analysis have rarely been directly confirmed in domestic cats, with most predictions being extrapolated from results in humans and mice.

To advance the field of basic veterinary medicine and support clinical veterinary practices not only for domestic cats, but also for various domestic and companion animals, it is crucial to accumulate a substantial amount of diverse basic data.

## Author Contributions

AA, KW and NS conceived and designed the study. AA and DK performed the analyses. AA and NS drafted the manuscript.

## Acknowledgements

This work was supported by a Grant-in-Aid for Scientific Research (KAKENHI) [Grant Number 21K06124] from the Japan Society for the Promotion of Science.

Computations were partially performed on the NIG supercomputer at ROIS National Institute of Genetics.

