## Supplementary material for "Possibilities of skin coat color-dependent risks and risk factors of squamous cell carcinoma and deafness of domestic cats inferred via RNA-seq data": RE_RE_Neko_ATWS_sup.docx

**Supporting Information**

**1. SCDEGs and results of KEGG pathway enrichment analysis of mesenteric lymph node cells from four different domestic short hair cats**

The estimation of SCDEGs and KEGG pathway enrichment analyses were performed using RNA-seq read data of mesenteric lymph node cells from four different domestic short hair cats (GSM4623155, GSM4623156, GSM4623157, and GSM4623158) on Gene Expression Omnibus (GEO). Here, the size factors (MORs) for these RNA-seq read data (Table S6a) were obtained 1.206467092, 0.840896415, 0.933185101, and 1.069045024, respectively, and [standard deviation of MOR over four cells] / [average MOR over four cells] was obtained 0.136652371. These results suggest that the size factor does not significantly affect the comparison of TPM values. Therefore, SCDEGs were chosen in a similar manner as the analysis of RNA-seq data of T, O, and W cells described in the text.

The SCDEG was defined as the gene that met the following conditions:

- The maximum expression levels obtained for 4 samples higher than *u*-times of the minimum value.
- The maximum expression levels obtained for 4 cells were equal to *v* or higher.

Here, $u=2$ and $v=$ [twice the median value of TPM values] were assumed to estimate SCDEGs.

To investigate the relationship between SCDEGs and the risk of squamous cell carcinoma, KEGG pathway enrichment analysis was conducted using g:Profiler (Table S6b). In this case, differently from T, O, and W cells, the following two squamous cell carcinoma related pathways, "pathways in cancer" and "Human papillomavirus (HPV) infection" were not the ones with significantly larger number of SCDEGs involved (Table S6b). This result may support that the obtained SCDEGs among three skins with different coat-colors and their squamous cell carcinoma related characteristics in the present study were not the nonspecific ones that emerged due to just only the variation among individuals.

**2. Ratio between d1KIT and normal KIT expressions estimated by maximum likelihood estimate (MLE)**

A simple model consisting of only two isoforms of KIT, normal KIT and d1KIT, without any bias in read distribution was considered. Afterward, MLE for the ratio [d1KIT expression]$/$[Normal KIT expression] is given as $=(r-1)/p-r$ with $p=n/N$ and $r=L/(L-l)$, where $n$ indicate the number of reads in KIT's 1st exon, $N$ indicate the mapped reads on all exons of KIT, $L$ indicate the length of all exons of KIT, and $l$ indicate the length of KIT's 1st exon (Pachter, 2011). Using the data of the number of read and the region length of each exon of KIT (Table S2.), the following results were obtained for T, O, and W cells, respectively:

p = 0.028842581, 0.027332262, and 0.016476662; r = 1.03539591, 1.03539591, and 1.03539591; and [d1KIT expression]$/$[Normal KIT expression] = 0.191814285, 0.259627163, and 1.112849284.

The expression level of d1KIT in T and O cells was quite smaller than those of normal KIT; however, the expression level of d1KIT in W cell seemed similar to that of normal KIT. Additionally, these results were consistent to the results shown in the Result section, where the ratio [TPM of d1KIT]$/$[TPM of normal KIT] were 0.179929164, 0.244270964, and 1.063338179 for T, O, and W cells, respectively.

**Table S1:** Number of reads and TPM of each gene in T, O, and W cells.

Table_S1.xlsx

**Table S2:** Number of reads and TPM of each exon in the KIT gene in T, O, and W cells.

Table_S2.xlsx

**Table S3:** List of SCDEGs and their TPMs in each cell, T, O, and W.

Table_S3.xlsx

**Table S4:** Squamous cell carcinomas types and the number of oncogenes and tumor suppressor genes in each squamous cell carcinoma.

Table_S4.xlsx

**Table S5**: Results of KEGG pathway enrichment analysis of SCDEGs using *u* = 2 and *v* = 2 (a) and *u* = 2.5 and *v* = 4.5 (b). Number of squamous cell carcinoma promoting and suppressor genes, which are strongly expressed in T (P_T_ and S_T_), O (P_O_ and S_O_), and W (P_W_ and S_W_) cells, and *P*-values of exact test of goodness-of-fit between promoting-suppressor genes of each case and the theoretical ratio using *u* = 2.5 and *v* = 4.5 (c). Here ALL indicate the theoretical ratio for all carcinomas and SCC indicate all squamous cell carcinomas classified in CancerMine (Table S4).

(a) Table_S5a.xlsx

(b) Table_S5b.xlsx

(c)

|  | T | O | W | ALL | SCC |
| --- | --- | --- | --- | --- | --- |
| Carcinoma promoting genes | P_T_ = 19 | P_O_ = 24 | P_W_ = 22 | 18994 | 904 |
| Carcinoma suppressor genes | S_T_ = 8 | S_o_ = 3 | S_W_ = 3 | 11295 | 584 |
| *P*-value (for ALL) | 0.4365 | 0.004415 | 0.006993 |  |  |
| *P*-value (for SCC) | 0.3328 | 0.002413 | 0.003887 |  |  |

**Table S6:** Number of reads and TPM of each gene in mesenteric lymph node cells of four different domestic short hair cats (a), and the result of the KEGG pathway enrichment analysis (b).

(a) Table_S6a.xlsx

(b) Table_S6b.xlsx

**Table S7:** List of pathways and roles of SCDEGs that are related to squamous cell carcinoma (top) and references (bottom).

Table_S7.xlsx

**Table S8:** List of genes that are related to melanoblast and melanocyte formation (top) and references (bottom).

Table_S8.xlsx

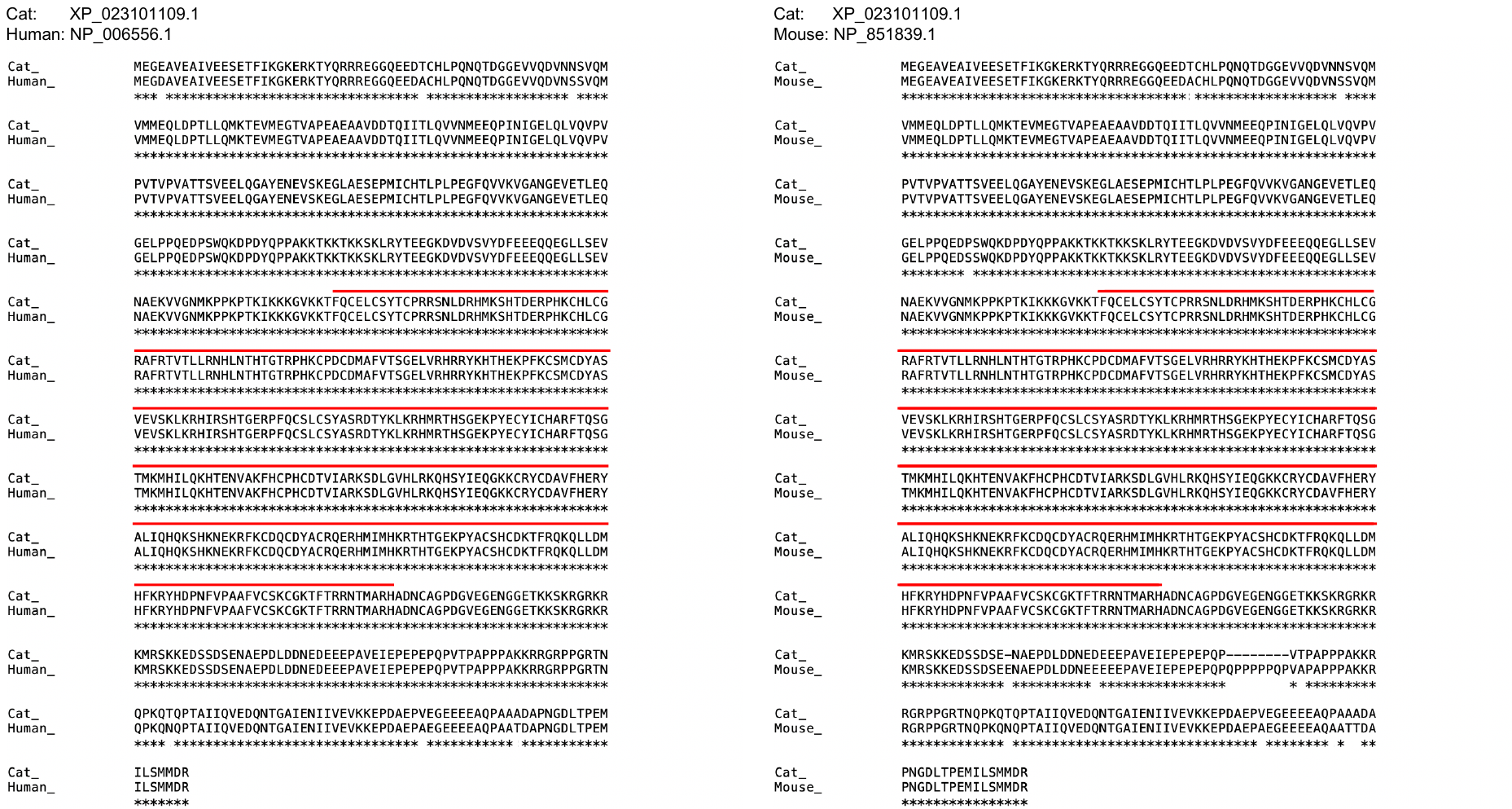

**Figure S1:** Comparison of amino acid sequences of CTCFs among human (NP_006556.1), mice (NP_851839.1), and domestic cat (XP_023101109.1). Sequence regions under red bars were DNA binding regions with 11 zinc-finger domains. Sequences above the asterisk represent homology between species. Among human, mice, and domestic cat, DNA binding region was completely conserved (a)(b). Between human and domestic cat, more than 99% of the entire CTCF protein sequence was conserved, with only 6 replacements found in 727 amino acid sequences (a).
